# A biosafety level-2 dose-dependent lethal mouse model of spotted fever rickettsiosis: *Rickettsia parkeri* Atlantic Rainforest-like isolate

**DOI:** 10.1101/488494

**Authors:** Andrés F. Londoño, Nicole L. Mendell, David H. Walker, Donald H. Bouyer

## Abstract

**Background:** The species of the *Rickettsia* genus is separated into four groups: the ancestral group, typhus group, transitional group and spotted fever group. *Rickettsia parkeri*, a spotted fever group *Rickettsia*, has been reported across the American continents as infecting several tick species and is associated with a relatively mild human disease characterized by eschar formation at the tick feeding site, fever, myalgia and rash. Currently several mouse models that provide a good approach to study the acute lethal disease caused by *Rickettsia*, but these models can only be performed in an animal biosafety level 3 laboratory. We present an alternative mouse model for acute lethal rickettsial disease, using *R. parkeri* and C3H/HeN mice, with the advantage that this model can be studied in an animal biosafety level 2 laboratory.

**Principal findings:** In the C3H/HeN mouse model, we determined that infection with 1 × 10^6^ and 1 × 10^7^ viable *R. parkeri* Atlantic Rainforest-like isolate produced dose-dependent severity, whereas infection with 1 × 10^8^ viable bacteria resulted in a lethal illness. The animals became moribund on day five or six post-infection. The lethal disease was characterized by ruffled fur, erythema, labored breathing, decreased activity, and hunched back, which began on day three post-infection (p.i.) and coincided with the peak bacterial loads. Significant splenomegaly (on days three and five p.i.), neutrophilia (on days three and five p.i.), and thrombocytopenia (on days one, three and five p.i.) were observed.

**Significance:** The greatest advantage of this inbred mouse model is the ability to investigate immunity and pathogenesis of rickettsiosis with all the tools available at biosafety level 2.

**Author summary:** *Rickettsia* is a bacterial genus that contains distinct species that are transmitted by arthropods. Many of these agents produce infection and disease in humans. The illness can range from very aggressive, such as Rocky Mountain spotted fever caused by *Rickettsia rickettsii*, to mild human disease characterized by eschar formation at the tick feeding site and less severe symptoms caused by *Rickettsia parkeri*. To study these diseases, animal models are invaluable, and mouse models offer the best advantages. Several mouse models are most useful for study of the acute lethal disease produced by these bacteria, providing the opportunity to test different treatments and vaccine candidates. However, work with these models requires an animal biosafety level 3 laboratory. In this report, we present an alternative mouse model to study acute lethal spotted fever group rickettsial disease with the advantage that experiments can be performed at biosafety level 2.

## Introduction

*Rickettsia*, a genus of Alphaproteobacteria that contains microorganisms transmitted by arthropods, is divided into four groups: the ancestral group, typhus group (e.g. *Rickettsia typhi*), transitional group (e.g. *R. australis*) and spotted fever group (e.g. *R. conorii* and *R. parkeri*) [1,2].

*Rickettsia parkeri*, a member of the SFG, has been reported across the American continents as infecting several tick species including the *Amblyomma maculatum* complex (*A. maculatum*, *A. triste*, *A. tigrinum*), *A. ovale*, *A. nodosum*, *A. parvitarsum* and *Dermacentor parumapertus* [3,4]. Distinct *R. parkeri* strains have been described in the American continents, including the Atlantic Rainforest strain isolated in Brazil [5]. The first isolate of *R. parkeri* in Colombia is most closely related to the Atlantic Rainforest strain from Brazil [6] and Black Gap strain from Texas [4]. *Rickettsia parkeri* has been associated with a relatively mild human disease characterized by eschar formation at the tick feeding site, fever, myalgia and rash [7–9].

Previously described mouse models provide a good approach to study the acute lethal disease produced by *Rickettsia*, such as *R. australis* infection of C57BL/6 or Balb/c mice, *R. conorii* infection of C3H/HeN mice and *R. typhi* infection of C3H/HeN mice [10–12]. The first one can be utilized to study a highly invasive model of rickettsial disease, because infection involves not only endothelial cells, but also perivascular cell types such as macrophages and can utilize gene knockout mice on the C57BL/6 background [10]. The second mouse model is the best available model for SFG rickettsial diseases, because the principal target cells are the vascular endothelium [11]. The third one is useful to study immunity and pathogenesis in typhus group rickettsial infection [12].

Analysis of the *R. parkeri* Atlantic rainforest-like (RpARFL) genome revealed several insertions, deletions and inversion zones in comparison to the North American *R. parkeri* Portsmouth strain; RpARFL contains bacterial conjugation genes including *traC*-, *traB*-, *traU*-and *traN*-like genes. The sequence of RpARFL has 1,348,030 bp that is 47,644 bp longer than the *R. parkeri* Portsmouth strain genome (unpublished). The aim of this study is to describe the clinical course, outcome, pathologic lesions, and anatomic distribution of rickettsiae in this new mouse model of SFG rickettsiosis.

## Materials and Methods

### Rickettsial quantification

A low-passage stock of RpARFL isolate grown in Vero cells was used for animal experiments [6]. This stock was passaged eight times in Vero cells and once in C3H/HeN mice to remove *Mycoplasma* sp. contamination. Rickettsial identity and *Mycoplasma sp*. removal were confirmed by full genome MINIon and Illumina sequencing. The infected Vero cells were harvested and then frozen in sucrose-phosphate-glutamate buffer solution (218 mM sucrose, 3.76 mM KH_2_PO_4_, 7.2 mM K_2_HPO_4_, and 3.9 mM glutamate). Rickettsial quantification was performed using a qPCR assay to measure the viable bacteria. Briefly, a six-well plate with confluent Vero cells was inoculated with 1 ml of RpARFL stock at a 1:100 dilution in 1% bovine calf serum (BCS) medium. The medium was aspirated from each well, and the wells were infected in triplicate or left as negative controls. The plate was centrifuged at 2,000 RCF for five minutes and incubated at 37°C for one hour. After the one hour rickettsial adsorption and entry into the cells, the plate was washed three times with warm (37°C) sterile PBS to remove extracellular rickettsiae and incubated again for one hour with 1 ml of 1% BCS growth medium with DNase (10U/µl) to degrade DNA of non-viable rickettsiae. The plate was washed three times followed by DNA extraction using the DNeasy Blood and Tissue Kit (Qiagen, Hercules, CA). For this purpose, 200 µl of PBS and 200 µl of the buffer AL were added to each well and incubated for 2 minutes. The contents of each well were collected in a clean tube and incubated with 20 µl of proteinase K at 56°C for 10 minutes. A positive control consisted of DNA from 10 µl of the original stock with 190 µl of PBS that was extracted in parallel. The DNA was further extracted following the manufacturer’s protocol. Primers for the single copy *gltA* gene, CS-5 (GAGAGAAAATTATATCCAAATGTTGAT) and CS-6 (AGGGTCTTCGTGCATTTCTT), were used for quantitative real-time PCR (qPCR) [13] with iTaq Universal SYBR Green Supermix (Bio-Rad, Hercules, CA), and the standard curve was prepared with dilutions (10^9^ to 10^1^ copies/µl) of a *R. conorii gltA* PCR fragment-containing plasmid. The concentration of the RpARFL stock used for this study was determined to be 3.72 × 10^9^ viable bacteria per ml. This value was utilized to calculate the doses of rickettsiae (1 × 10^8^, 1 × 10^7^ or 1 × 10^6^ bacteria) in a total volume of 200 µl.

### Mice

Eight week old male C3H/HeN mice from Charles River Laboratories, Inc. (Houston, TX) were used in this study. The experiments were performed in an animal biosafety level 3 facility, under specific pathogen-free conditions. All animal work was approved by the Institutional Animal Care and Use Committee (protocol # 9007082) of the University of Texas Medical Branch-Galveston prior to permission to use *R. parkeri* at BSL-2, and mice were used according to the guidelines in the Guide for the Care and Use of Laboratory Animals and comply with the USDA Animal Welfare Act (Public Law 89-544), the Health Research Extension Act of 1985 (Public Law 99-158), the Public Health Service Policy on Humane Care and Use of Laboratory Animals, and the NAS Guide for the Care and Use of Laboratory Animals (ISBN-13).

### Preliminary dose range experiment

To determine the susceptibility to and lethal dose of RpARFL in a murine model, a dose range-finding experiment was undertaken. Groups of mice (n=3/group) were inoculated intravenously (I.V.) with 1 × 10^6^, 1 × 10^7^ or 1 × 10^8^ viable bacteria or PBS (200 µl). Animals were monitored daily for signs of illness, rectal temperature and body weight.

### Infection assays

To characterize the kinetics of the lethal murine model of infection with RpARFL, three groups of animals (n=6/group) were infected with 200 µl of the highest dose (1 × 10^8^) I.V, and one group was utilized as a control (200 µl of PBS I.V). Due to the kinetics of the high dose infection, which resulted in morbidity by day six p.i., infected mice were sacrificed on days one, three and five post-infection (p.i.), and the animals from the control group (n = 6), on day six. As in the first study, the animals were monitored daily. The spleen, kidneys, liver, lungs, heart and brain were collected from each animal to determine bacterial loads and evaluate histopathologic changes. We collected blood samples in BD microtainer tubes with and without K_2_EDTA (Becton Dickinson, Franklin Lakes, NJ) to analyze the blood cell counts, determine bacterial loads, and assess the serologic response.

### Hematology and serology

Blood samples were collected in two microtainer tubes, with and without K_2_EDTA, from each animal at the time of sacrifice. The complete blood cell counts were measured with a HemaVet 950FS apparatus (Drew Scientific, Miami Lakes, FL). Indirect immunofluorescence assay (IFA) was performed to measure IgG and IgM antibodies to RpARFL using acetone-permeabilized RpARFL-infected Vero cell-coated slides. The slides were immersed in phosphate buffered saline (PBS) for 10 minutes at room temperature, transferred to blocking solution (PBS with 1% bovine serum albumin [BSA] and 0.01% sodium azide), and incubated for 15 minutes. Sera were diluted in a series of two-fold dilutions starting at 1:64 in a solution of PBS with 1% BSA and 0.1% Tween-20. Experimental samples of serum as well as one positive control and one negative control serum per slide were added to the wells and incubated at 37°C for 30 minutes in a humidified chamber. For IgM titer determination, IgG binding was blocked by treating the serum with IgM Pretreatment Diluent (Focus Diagnostics, California, USA) prior to assaying. The slides were washed two times for 10 minutes in PBS containing 0.1% Tween-20. Secondary antibody, DyLight 488-conjugated anti-mouse IgG (1:15,000 dilution, Jackson Immunoresearch, West Grove, PA) or FITC-conjugated anti-mouse IgM antibody, mu chain specific (1:500 dilution, Vector Laboratories, Burlingame, CA), was added to the wells and incubated for 30 minutes in a humidified chamber. Finally, the slides were washed twice as before with the final wash containing 1% Evans blue solution, mounted with DAPI fluoromount-G (SouthernBiotech, Birmingham, AL), and coverslipped. Slides were observed under a fluorescence microscope at 400X magnification (Olympus Scientific, Waltham, MA).

### Measurement of rickettsial loads by real-time PCR

DNA was extracted from tissue and blood using a Qiagen DNeasy Blood and Tissue Kit (Qiagen, Hercules, CA), following the manufacturer’s protocol. The bacterial loads were determined by qPCR using the primers CS-5 and CS-6 with the probe (FAM - CATTGTGCCATCCAGCCTACGGT), and iQ Supermix (Bio-Rad, Hercules, CA) [13,14]. The standard curve was determined as described above in a qPCR assay to measure the viable bacteria. Samples were normalized using tissue weight or blood volume, and the concentration of rickettsiae is expressed as gene copies per milligram of tissue or milliliter of blood.

### Histology and Immunohistochemistry

The tissues were fixed with 10% neutral buffered formalin for two weeks and embedded in paraffin. The samples were sectioned at 5 µM thickness and stained with hematoxylin and eosin for histological analysis or processed for immunohistochemical (IHC) staining of rickettsial antigen [15].

The sections were incubated at 54°C overnight, deparaffinized and hydrated. After that, they were blocked with Avidin/Biotin Blocking Kit (Life Technologies, Frederick, MD) and treated with proteinase K (Dako, Carpinteria, CA), for antigen retrieval. The sections were incubated with polyclonal rabbit ant-*R. conorii* antibody (1:300 dilution, produced in-house) at room temperature for one hour, followed by biotinylated secondary anti-rabbit IgG (1:200 dilution, Vector Laboratories, Burlingame, CA), streptavidin-AP (1:200, Vector Laboratories, Burlingame, CA) for 30 minutes each and Fast Red (Dako, Carpinteria, CA) for 5 minutes. The sections were washed twice with Tris-buffered saline containing 0.05% Tween-20. Slides were counterstained with hematoxylin, dehydrated, mounted with Permount and examined with an Olympus BX51 microscope (Olympus Scientific, Waltham, MA).

### Statistical analysis

Data were analyzed using GraphPad Prism software utilizing a one-way ANOVA with Tukey’s post-test. P-values of <0.05 were considered to indicate statistically significant differences in each analysis.

## Results

### Susceptibility and lethal dose of RpARFL in a murine model

In C3H/HeN mice, we observed that infection with 1 × 10^6^ (low dose) and 1 × 10^7^ (mid dose) of viable *R. parkeri* ARFL produced dose-dependent severity of illness. The animals which received the lowest concentration doses began to show signs of illness on day three p.i. as evidenced by onset of weight-loss. Animals in the lowest dose group presented with piloerection on days three and four p.i.; whereas piloerection on day three, and piloerection, erythema, and hunched back were observed on day four in the mid dose group, which diminished to piloerection alone by five days p.i. By days five and six p.i., the mice of the low- and mid-dose groups, respectively, did not have observable signs of illness through the experiment’s end (day 14). The control group did not present with any sign of illness during the experimental observation period. Infection with 1 × 10^8^ (high dose) of viable bacteria resulted in a lethal illness. The animals became moribund on day six p.i., when the mice had lost 14.6% of their initial body weight and developed hypothermia (Fig. 1A and B).

**Fig 1.**
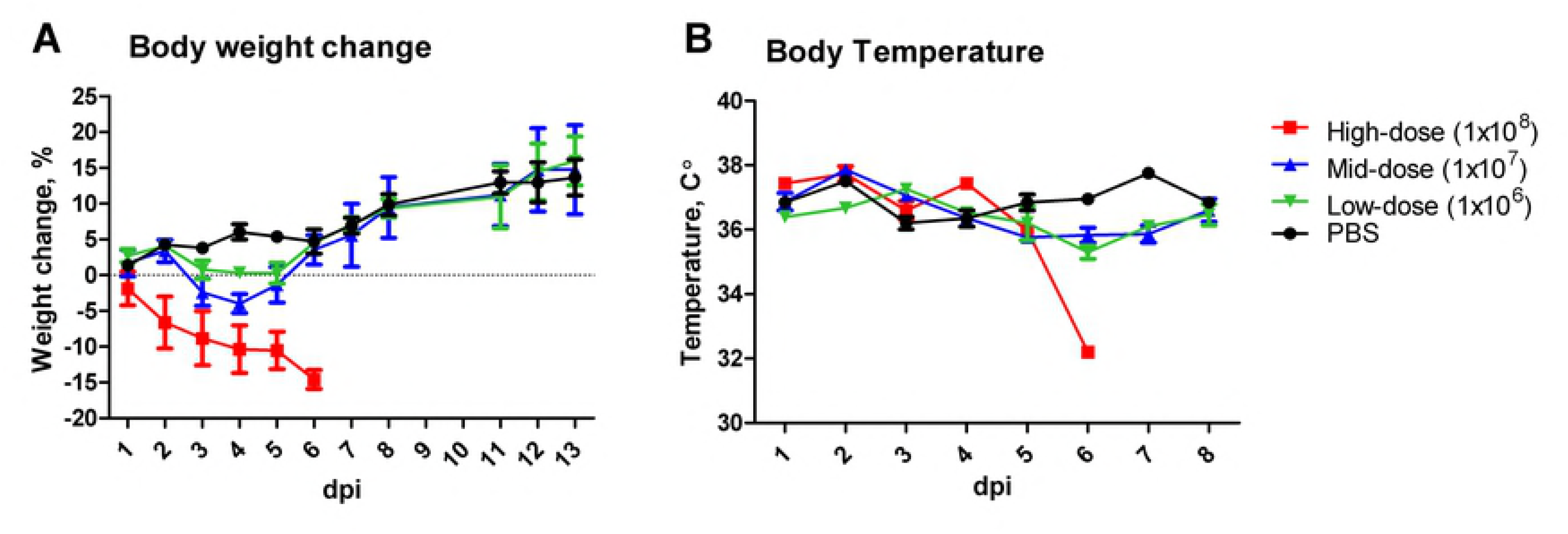
Kinetics of body weight (**A**) and temperature changes (**B**) in C3H/HeN mice infected intravenously with high-(1 × 10^8^, square), mid-(1 × 10^7^, triangle) and low-(1 × 10^6^, inverted triangle) doses of viable RpARFL or PBS inoculated controls (circle).

### Kinetics of signs of illness in the lethal murine model of infection with RpARFL

The disease was characterized by ruffled fur, erythema, labored breathing, decreased activity, and hunched back, which began on day three p.i., and the severity of these signs intensified with time. The decreased body weight and body temperature observed in the dose-range study were replicated (Fig. 2A and B). Significant splenomegaly was observed on days three and five when compared with the control group (Fig. 2C).

**Fig 2.**
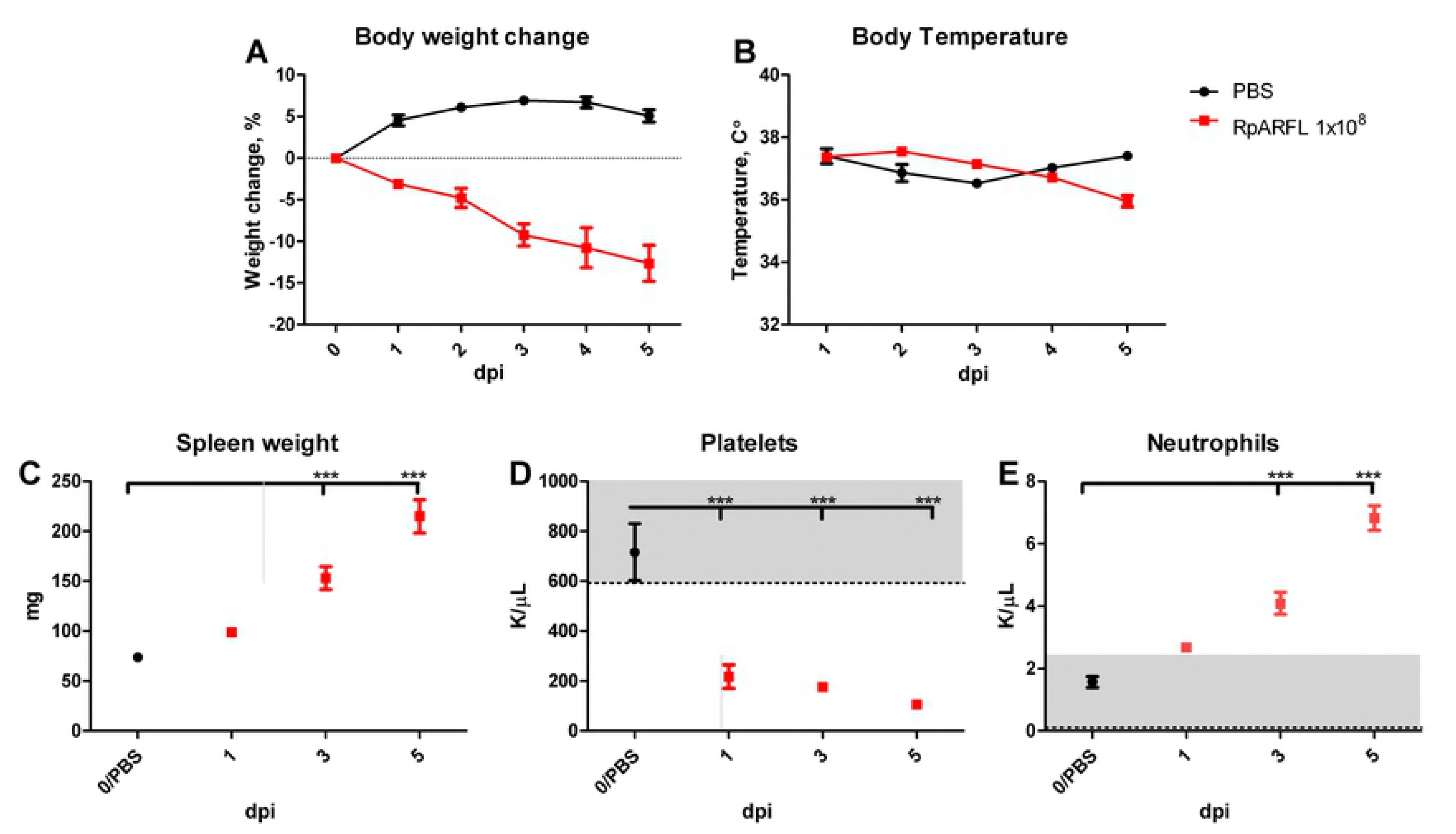
Kinetics of changes in weight (**A**), body temperature (**B**), and spleen weight (**C**) of mice infected intravenously with 1 × 10^8^ viable RpARFL organisms and uninfected controls. Mean platelet (**D**) and mean neutrophil concentrations (**E**) of mice infected intravenously with 1 × 10^8^ viable RpARFL organisms on days 1, 3 and 5 p.i. and uninfected controls measured with a HEMAVET 950FS apparatus.

### Hematology and serology

The most marked findings in the complete blood cell counts of lethally infected animals were significant thrombocytopenia on days one, three and five p.i. that gradually progressed in severity each day and neutrophilia on days three and five p.i. (Fig. 2D and 2E). The animals infected with the low- and mid-doses developed a strong antibody response measured by IFA on day 14 p.i. The IgM reciprocal endpoint titers were between 1,024 and 4,096 for both groups and the IgG titers between 8,192 and 16,384 for the animals that received the low dose, and 16,384 and 32,768 for the animals that received the mid dose. IgM antibody was detectable on day three in the animals that received the high dose with reciprocal endpoint titers between 256 and 512, and between 512 and 1,024 on day five. In contrast, IgG antibody was present in only two of the six mice on day five at the cutoff reciprocal titer of 64.

### Measurement of rickettsial loads by real-time PCR

All tissues of RpARFL-infected mice assayed contained rickettsial DNA. In the spleen, lung and liver, qPCR analysis revealed that the peak bacterial loads occurred on day three p.i., with statistical significance, and decreased on day five p.i. (Fig. 3A, B and C). In the heart and kidney, the bacterial loads appeared to peak on day three, but without differences among the days of infection reaching statistical significance (Fig. 3D and E). Infection was observed in the brain on days three and five p.i., with no statistical significance between the time points (Fig. 3F). In the blood, the peak bacterial load was observed on day five p.i. (Fig. 3G).

**Fig 3.**
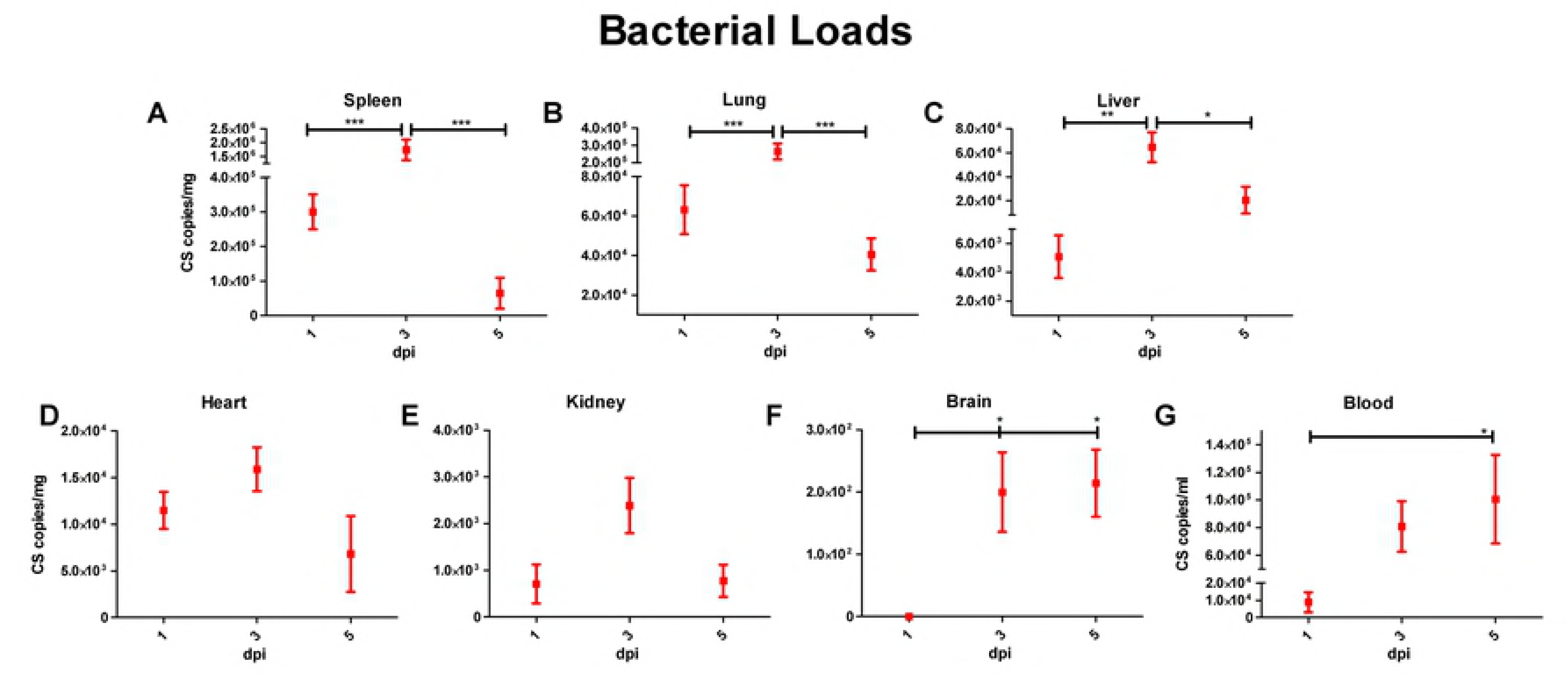
Measurement of rickettsial loads in spleen (**A**), lung (**B**), liver (**C**), heart (**D**), kidney (**E**), brain (**F**), and blood (**G**) expressed as copies of citrate synthase (CS) gene per milligram of tissue or per milliliter of blood on days 1, 3 and 5 p.i. of mice infected with 1 × 10^8^ viable RpARFL organisms.

### Histology and Immunohistochemistry

No significant pathologic findings were detected in the tissues of mice infected with low- and mid-dose of RpARFL or the uninfected control mice. Meningitis was observed in the brains of animals infected with the high dose of RpARFL on days three and five p.i. (Fig. 4A and B). Pathology in the heart was characterized by mural and valvular endocarditis and perivascular interstitial cellular infiltration beginning on day three (Fig. 5A). The kidney showed inflammatory mononuclear cellular infiltration between the renal tubules and intertubular capillaries (Fig. 6A). Interstitial pneumonia was observed beginning on day three p.i. (Fig. 7A). Cellular infiltration in liver was characterized by a high ratio of polymorphonuclear cells to mononuclear cells on day three p.i., and the inverse was observed on day five p.i., fewer polymorphonuclear cells and an increase in mononuclear cells (Fig. 8A and B). RpARFL organisms were associated with vessels of the microcirculation in areas of pathological damage in the brain, lung, kidney and heart by immunohistochemical staining (Fig. 4C, D, 5B, 6B and 7B). In the liver tissue RpARFL was observed in hepatocytes and mononuclear cells in addition to endothelial cells (Fig. 9A, B, C and D).

**Fig 4.**
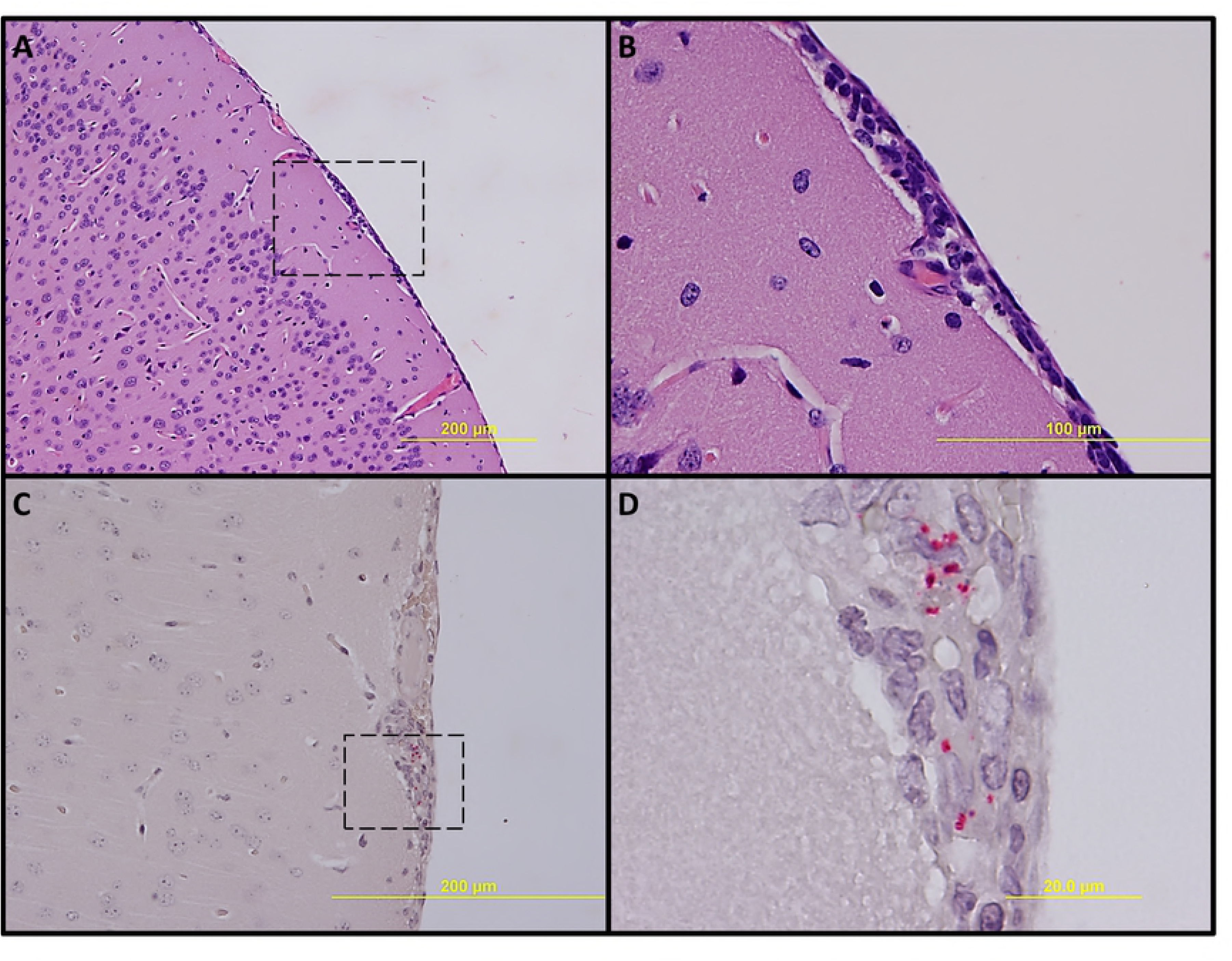
Pathologic lesions and IHC demonstration of SFG rickettsiae in brain. Meningitis on day 3 p.i. (**A**, 100X) and inset (**B**, 400X). Meningitis with presence of RpARFL (red) (**C**, 200X) and inset (**D**, 1,000X) on day 5 p.i.

**Fig 5.**
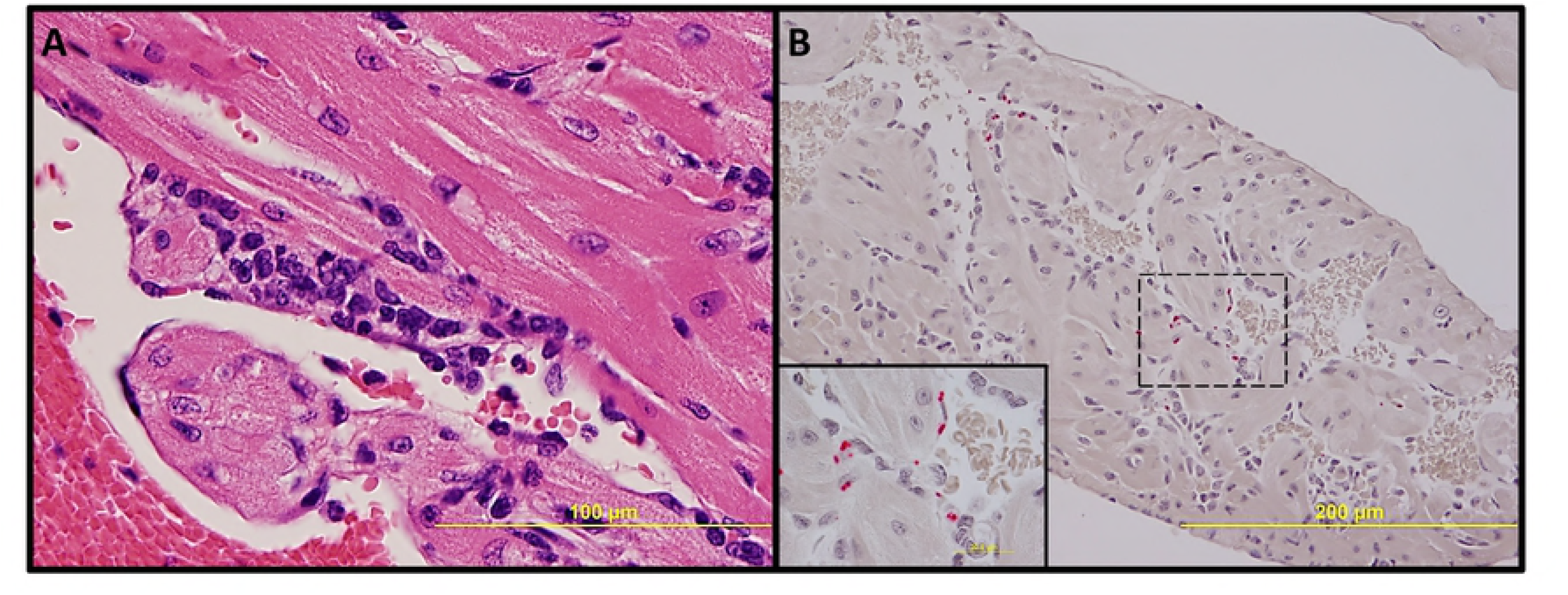
Pathologic lesions and IHC demonstration of SFG rickettsiae in heart. Endocarditis and perivascular interstitial inflammation on day 3 p.i. (**A**, 400X). Endothelial presence of RpARFL (red) (**B**, 200x) and inset (1,000X) on day 3 p.i.

**Fig 6.**
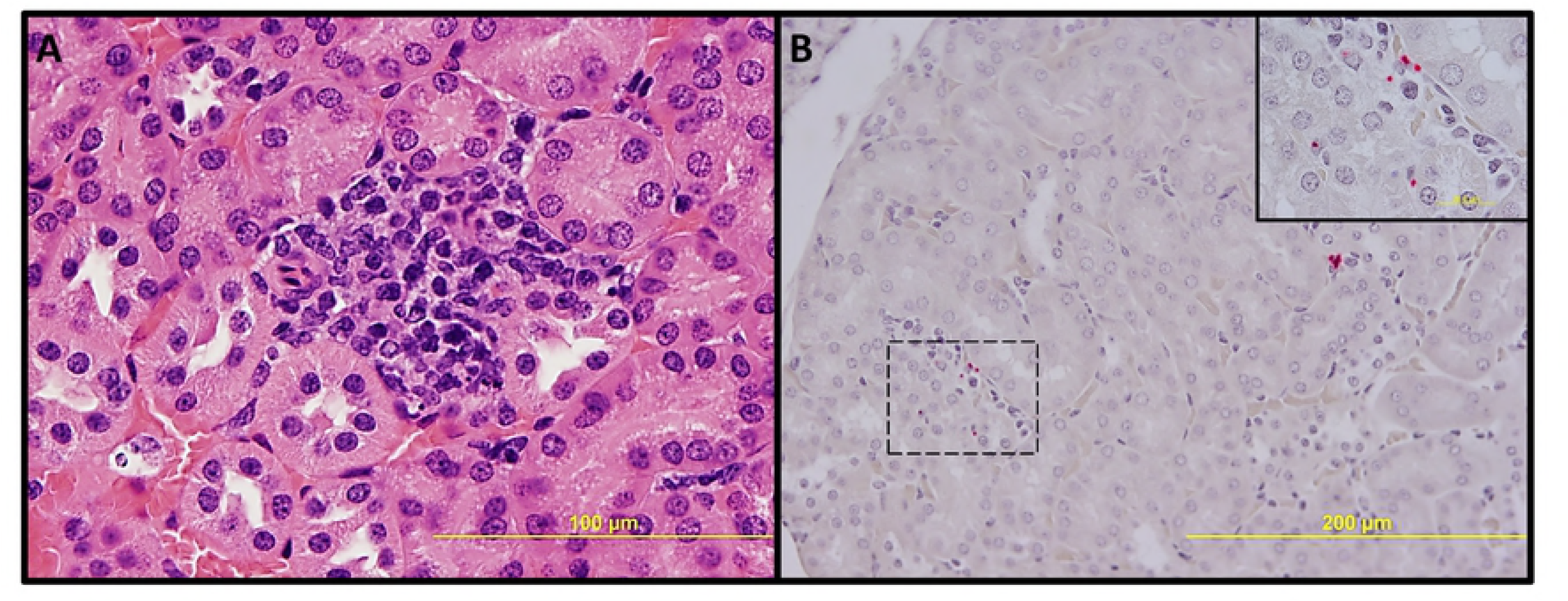
Pathologic lesions and IHC demonstration of SFG rickettsiae in kidney. Inflammation with interstitial mononuclear cellular infiltration on day 3 p.i. (**A**, 400X). Endothelial presence of RpARFL (red) (**B**, 200X) and inset (1,000X) on day 3 p.i.

**Fig 7.**
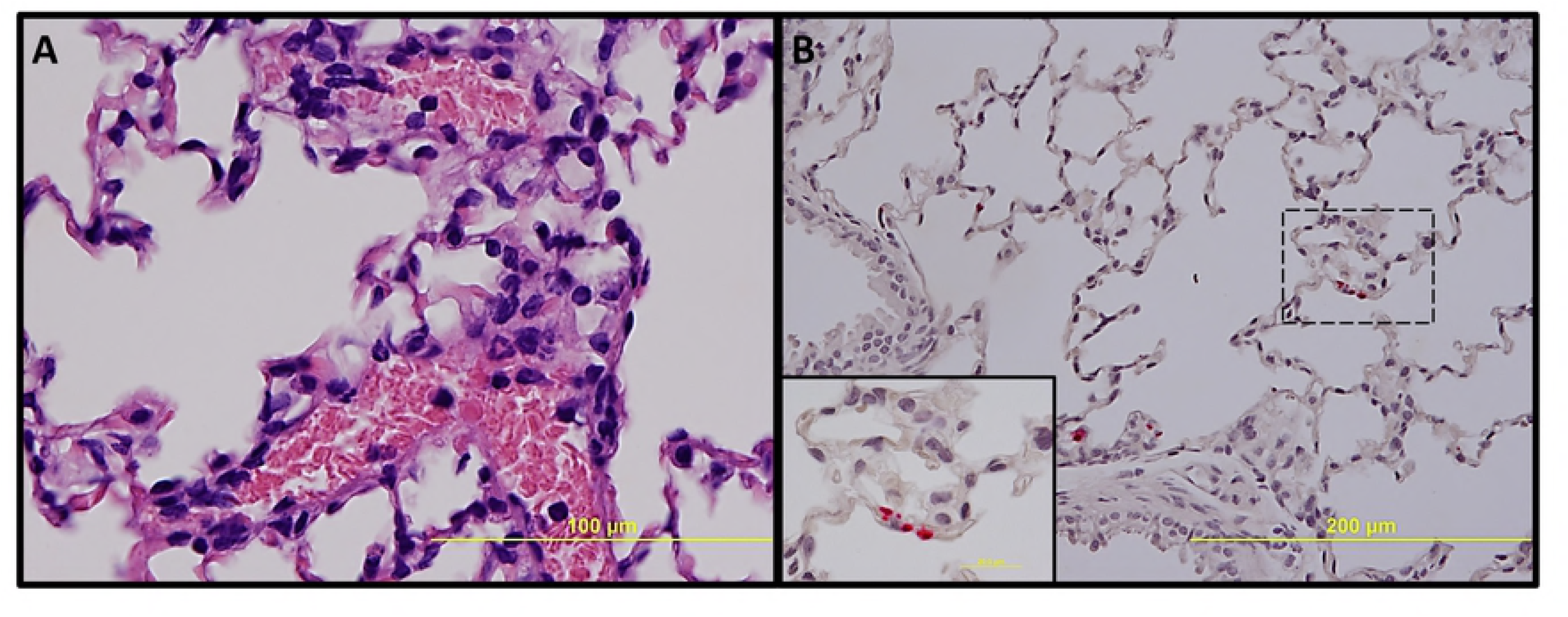
Pathologic lesions and IHC demonstration of SFG rickettsiae in lung. Interstitial pneumonia on day 3 p.i. (**A**, 400X). Endothelial presence of RpARFL (red) in alveolar capillaries (**B**, 200X) and inset (1,000X) on day 3 p.i.

**Fig 8.**
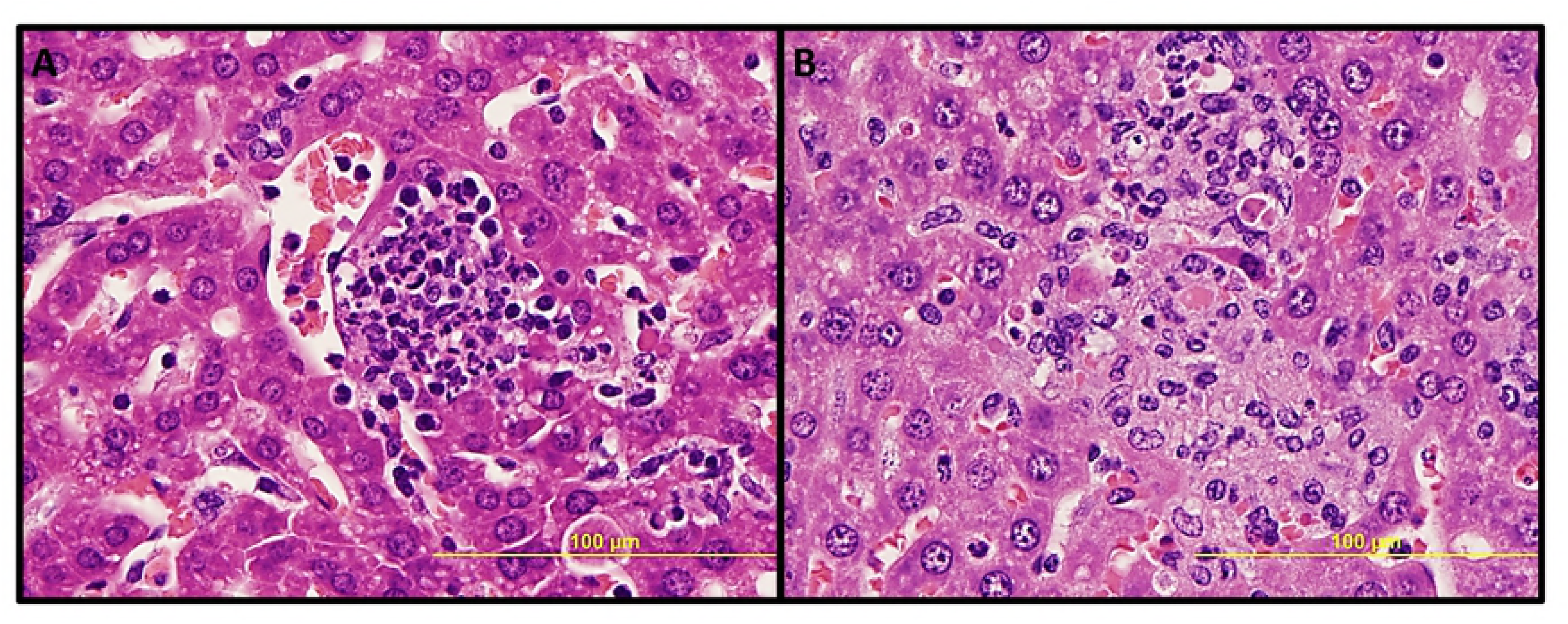
Pathologic lesions in liver. Focal inflammation with polymorphonuclear cells predominance on day 3 p.i. (**A**, 400X), and with mononuclear cells predominance and hepatocyte apoptosis on day 5 p.i. (**B**, 400X).

**Fig 9.**
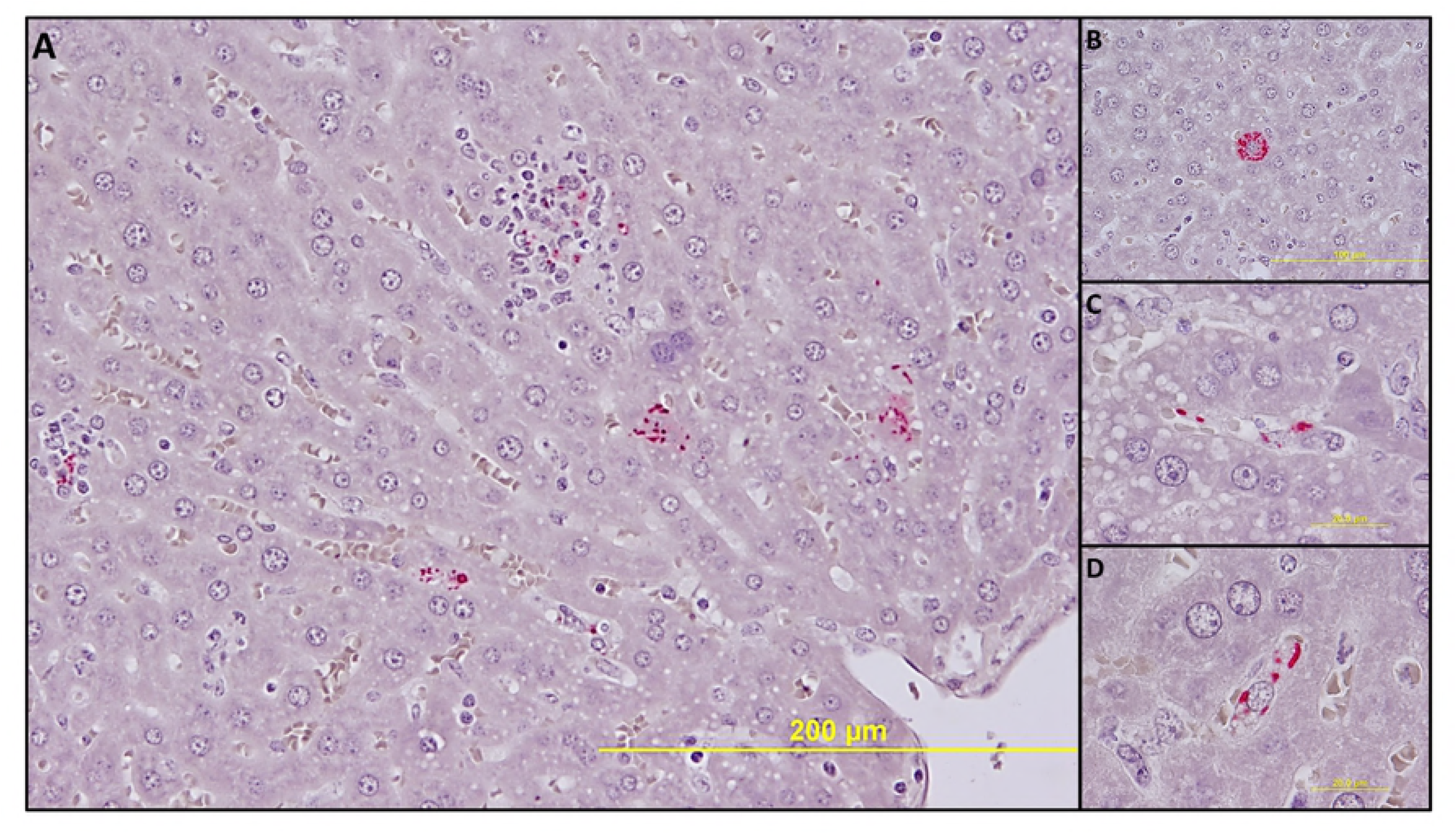
IHC demonstration of SFG rickettsiae in liver. Rickettsiae (red) in multiple types of infected cells (**A**, 200X). In a hepatocyte (**B**, 400X), in endothelial cells (**C**, 1,000X) and in mononuclear cells (**D**, 1,000X) on day 3 p.i.

## Discussion

This experimental infection provides an alternative animal model in mice for rickettsial disease with the advantage that this model can be studied in an animal biosafety level 2 (ABSL-2) laboratory. Three rickettsial mouse models, *R. australis* infection of C57BL/6 or Balb/c mice, *R. conorii* infection of C3H/HeN mice and *R. typhi* infection of C3H/HeN mice, can only be performed in an ABSL-3 laboratory [10–12].

Infection of C3H/HeN mice with low- and mid-dose RpARFL resulted in dose-dependent severity, whereas infection with the high dose produced a lethal illness. The animals became moribund by day five or six p.i. The lethal disease was characterized by ruffled fur, erythema, labored breathing, decreased activity, and hunched back, which began on day three p.i. and coincided with the peak bacterial loads in some tissues (Fig. 4). Other significant observations included splenomegaly (on days three and five p.i.), neutrophilia (on days three and five p.i.), and thrombocytopenia (on days one, three and five p.i.). Two previous studies also reported that *R. parkeri* causes dose-dependent severity in mice. In one study, the authors infected C3H/HeN mice with 5.5 × 10^6^ of the *R. parkeri* Portsmouth strain, and the animals sacrificed on day seven did not develop any clinical sign of illness. The mice displayed mild to moderate splenomegaly, and *R. parkeri* DNA was detected in heart and lung of only 50% of the animals [16]. In the other study, 1.2 × 10^7^ of *R. parkeri* Maculatum 20 strain caused lethargy, mild hunched posture, ruffled fur and decreased activity with onset around days 6-7 p.i. These clinical manifestations persisted for 2-3 days. Bacterial DNA was detected in lung, spleen, liver and brain on day six p.i. [15]. The data from these two reports of C3H/HeN mice infected by intravenous inoculation of *R. parkeri* are similar to our low- and mid-dose infections, respectively.

A study of infection of guinea pigs (*Cavia porcellus*) by intraperitoneal inoculation of 1.9 × 10^8^ of *R. parkeri* Black Gap strain resulted in a nonlethal illness, manifested in most animals by mild fever and mild to moderate swelling and erythema of the scrotum [4]. This strain has a close genetic identity with the Brazilian Atlantic rainforest strain [4] and the Colombian RpARFL isolate [6]. Furthermore, subclinical infection was observed in an additional study that utilized the natural mode of infection of guinea pigs by tick transmission, using *A. ovale* nymphs infected with the Brazilian Atlantic rainforest strain [17].

In conclusion, we have described an animal model using C3H/HeN mice infected by intravenous inoculation with a 1 × 10^8^ dose of RpARFL, which provides an opportunity to study acute lethal disease produced by SFG rickettsiae characterized by infection of endothelial cells in the brain, heart, lung and kidney, and endothelial cells, hepatocytes and mononuclear cells in liver. Clinical case reports of rickettsial infection on occasion incorrectly state that the patients had hepatic failure without clinical evidence such as increased levels of serum ammonia or decreased hepatic synthesis of proteins. Hepatic injury manifest as moderately increased serum concentration of alanine aminotransferase and aspartate aminotransferase occur frequently in spotted fever rickettsioses, and hyperbilirubinemia is observed in the most severe cases of Rocky Mountain spotted fever. However, histopathologic studies have not detected severe hepatic necrosis. This mouse model with prominent infection of a modest number of hepatocytes offers an opportunity to investigate the pathophysiology of focal hepatic injury in SFG rickettsial infection.

The greatest advantage of this inbred mouse model is the ability to investigate immunity and pathogenesis of rickettsiosis with all the tools available at biosafety level 2.

## Acknowledgement

The authors wish to express gratitude to Tais Saito for immunohistochemistry advice. Juan Olano and Bi-Hung Peng for kindly providing primary anti-SFG *Rickettsia* antibody, and Guang Xu for technical assistance. We also express our thanks to Colciencias for its support with the PhD scholarship for Dr. Londoño, through the official announcement 511/2010.

